# “Embryonic piRNAs target horizontally transferred vertebrate transposons in assassin bugs”

**DOI:** 10.1101/2024.01.22.576481

**Authors:** Tarcísio Fontenele Brito, Maira Arruda Cardoso, Nazerke Atinbayeva, Ingrid Alexandre de Abreu Brito, Lucas Amaro da Costa, Nicola Iovino, Attilio Pane

**Affiliations:** Instituto de Ciências Biomédicas, Universidade Federal do Rio de Janeiro, 21941-902 Rio de Janeiro, Brasil; 2 Instituto Nacional de Ciência e Tecnologia em Entomologia Molecular, Brasil; Department of Chromatin Regulation, Max Planck Institute of Immunobiology and Epigenetics, D-79108 Freiburg, Germany; Albert-Ludwigs-Universität Freiburg, Fahnenbergplatz, 79085 Freiburg im Breisgau, Germany

**Author notes:** Corresponding author (AP).

**Keywords:** *Rhodnius*, HTT, piRNA pathway, transposon, siRNA

## Abstract

Piwi proteins and the associated Piwi-interacting RNAs (piRNAs) coordinate a surveillance system that protects the animal genome from DNA damage induced by transposable element (TE) mobilization. While the pathway has been described in detail in the fruit fly *Drosophila melanogaster,* much less is known in more basal insects. Here, we investigated the adaptation of the piRNA pathway to horizontally transferred transposons (HTTs) in the assassin bug *Rhodnius prolixus*, a primary vector of Chagas disease. *Rhodnius* acquired specific classes of HTTs by feeding on bats, opossums and squirrel monkeys. By analyzing the temporal dynamics of piRNA cluster expression and piRNA production during critical stages of *Rhodnius* development, we show that peak levels of ∼28 nt long piRNAs correlate with reduced HTT and resident TE expression primarily during embryogenesis. Strikingly, while resident TEs piRNAs seem to engage in a typical ping-pong amplification mechanism, sense and antisense HTT piRNAs instead overlap by ∼20 nt or do not display ping-pong signatures. These features are explained at least in part by the low number of HTT copies inserted into the piRNA clusters and might point to a non-canonical mechanism of biogenesis. Our data reveal that the piRNA, but not the siRNA pathway, responded to HTTs that were recently transferred from vertebrate tetrapods to a hematophagous insect of medical relevance.

## Introduction

Transposable elements (TEs) are selfish genetic elements that can mobilize and insert in new positions of eukaryotic genomes. These sequences often comprise a substantial proportion of the host genome reaching more than half of the human genome and 90% of the maize genome [1–3]. Although TEs are usually passed on to the next generations via vertical transmission, a growing body of evidence is showing that horizontal transposons transfer (HTT) is more widespread than previously appreciated and occurs not only between closely related species, but also across distant phyla [4–6]. One of the most remarkable examples is represented by the horizontal transfer of DNA transposons between the triatomine insect *Rhodnius prolixus* and vertebrates, which occurred within the past 50 millions years [7]. *Rhodnius* is a hematophagous insect species and a primary vector of *Trypanosoma cruzi*, the causative agent of Chagas disease. Transposons of the *SPACE INVADERS* (*SPIN*), *OposCharlie1* (*OC1*), *hAT1* and *ExtraTerrestrial* (*ET*) in the genome of this insect were found to share up to ∼98.1% sequence identity with those described in opossums, squirrel monkeys and bats of South America [7]. *Rhodnius* likely acquired these HTTs because of its blood-feeding habit and the broad range of vertebrate hosts. The same habit is responsible for the transmission of *T. cruzi* to humans and underlies the etiology of Chagas disease. *Rhodnius* also harbors other promiscuous transposons like *RTE-X*, *Gypsy, Helitron, I, Maverick* and *Mariner*, with the latter being the predominant family in the *Rhodnius* genome [8]. These TEs are often transmitted horizontally across the animal kingdom and even between animals and plants [9]. How cells respond to newly invading transposons has been investigated in the fruit fly *Drosophila melanogaster*. The best characterized example of HTT involves the horizontal transfer of P-elements from *Drosophila willistoni* to *Drosophila melanogaster* and their subsequent spreading across natural *D. melanogaster* populations over the second half of the 20th century [10]. P-element invasion of a naive genome was soon associated with a class of developmental defects collectively known as hybrid dysgenesis [11,12]. Hybrid dysgenesis leads to sterility but only manifests itself in the progeny that paternally inherits the P-element. The discovery of the piRNA pathway and its crucial role as a defense system against transposon-induced DNA damage allowed to shed light on the molecular underpinnings of hybrid dysgenesis [13]. Piwi-interacting RNAs or piRNAs are a class of small non-coding RNAs ranging in size from 18 to 30 nucleotides that are typically found in complex with PIWI proteins [14–16]. The pathway has been best described in *Drosophila melanogaster* and a few other holometabolous insect species [17]. In *Drosophila* somatic cells, piRNA precursor transcripts originate from regions of the genome harboring a variety of transposon remnants known as piRNA clusters [18,19]. These piRNA precursors are processed in cytoplasmic organelles known as Yb-bodies to generate mature mostly antisense piRNAs [20]. In turn, antisense piRNAs are loaded into the Piwi protein that translocates to the nucleus and employs them as guides to identify and silence actively transcribed transposons in the genome [21,22]. Among the primary targets of the somatic pathway are the *Zam*, *Gypsy* and *Idefix* transposons [19,23,24]. The transcription of the germline piRNA clusters instead occurs from both genomic strands (i.e. dual-strand piRNA clusters) [19] and is regulated by a dedicated machinery [25–28]. Cluster transcripts produce primary antisense piRNAs as in the somatic cells [29]. However, a fraction of these RNAs promotes a feedforward amplification loop known as the “ping-pong” cycle, that is regulated by Aubergine (Aub) and Argonaute3 (Ago3) [18,19] . The loop starts when Aub bound to primary antisense piRNAs elicits the formation of sense piRNAs by cleaving the transcripts of active transposons. In turn, sense piRNAs are bound by Ago3, which can now recognize cluster transcripts to produce more antisense piRNAs. The slicing activities of Aub and Ago3 leave a typical hallmark in the secondary piRNA complement known as “ping-pong signature”, whereby sense and antisense piRNAs overlap by 10 nt at their 5’ end. Also, because antisense piRNAs display a 5’ Uracil bias (1U), the sense piRNA mates are enriched with an Adenine residue at the 10th position (10A). Antisense piRNAs associate with Piwi also in the germline and guide the protein to the nucleus.The ping-pong cycle is also involved in the production of phased primary piRNAs [30]. Once a piRNA precursor is cleaved by Ago3 or Aub to generate a secondary piRNA, the remaining sequence immediately after the 3’ end of the cleavage site is successively clipped and trimmed. The mechanism is widely conserved among animal species, increasing not only the number but also the diversity of piRNA sequences [30,31]. In addition to the piRNA pathway, endogenous ∼22 nt long siRNAs (siRNAs) have also been implicated in transposon downregulation in *Drosophila* [32,33]. siRNAs are generated through the cleavage double-strand RNA molecules by the ribonuclease Dicer2 (Dcr2) and loaded into Argonaute2 (Ago2) to form the RISC complex [34,35]. Transposon-related siRNAs can originate from piRNA clusters or from transposable elements [32]. In *Drosophila*, the siRNA pathway downregulates certain TEs in the somatic cells, while the piRNA pathway is crucial in germline tissues.

Since the discovery of horizontally transmitted P-elements in *Drosophila* species, a wealth of studies have reported the exchange of transposable elements between non-hybridizing insect species [5,12,36]. The piRNA pathway has been proposed to coordinate an adaptive defense mechanism against newly invading transposons [37]. A naive genome would not express piRNAs against a new TE, which can therefore insert in several positions in the genome. However, when a transposon copy eventually integrates in a piRNA cluster, it becomes a source of piRNAs that guide the PIWIs to silence in *trans* the active elements dispersed in the genome. The observation that piRNAs are maternally loaded in the developing eggs also pointed out a mechanism to explain the hybrid dysgenesis syndrome [13]. The progeny of a cross between a female fly bearing a P-element and a naive male will develop normally because it inherits both the TE and the cognate piRNA set. In contrast, the reciprocal cross will prove sterile because the P-element is transmitted by the father to a progeny that lacks the appropriate piRNA set. Aside from the Drosophilids however, much less is known about the mechanisms that allow insect species to repress newly invading transposons. The piRNA pathway is evolutionarily conserved in animals, but remarkable differences can be observed even when the investigation is restricted to species of the same genus [17,38,39]. For instance, it was recently shown that *D. eugracilis*, which diverged only 10 million years ago from *D. melanogaster*, neither uses a ping-pong amplification mechanism nor it expresses the critical YB protein [38]. The expression patterns and size distribution of the piRNAs also varies when *D. melanogaster* is compared to other holometabolous insects like *Tribolium casteneum*, *Bombyx mori* and *Apis mellifera* [40] [41–43]. In hemimetabolous insects the characterization of the pathway is still in its infancy, but it was shown that *Blattella germanica* expresses ∼28 nt piRNAs during embryogenesis as well as stage-specific piRNA pools during post embryonic development [41]. The investigation of the piRNA pathway in more basal insects therefore might reveal novel mechanisms and functions and shed light on the arms race between transposons and related defense systems.

*Rhodnius prolixus* is a hemimetabolous insect belonging to the Triatomine subfamily and undergoes 5 nymph stages during development before turning into a fertile winged adult [44]. Blood feeding is not only necessary for ecdysis, but also to trigger oogenesis in adult females. It was estimated that between 19% and 23% of the *Rhodnius* genome is composed of transposon sequences with the *Mariner* family accounting for a large fraction of the mobilome [8,45,46]. The genome of this hemipteran insect harbors four *PIWI* genes and we have previously shown that *Rp-piwi2*, *Rp-piwi3* and *Rp-ago3*, but not *Rp-piwi1*, are expressed in ovaries and are necessary for female adult fertility [47]. Furthermore, our transcriptomic analyses revealed that orthologs of the *Drosophila* piRNA and siRNA pathways components are also expressed in *Rhodnius* ovaries [48]. In this study, we characterize the temporal dynamics of the piRNAs and their source *loci* in the *Rhodnius* genome. Importantly, we provide the first evidence of the ability of the piRNA pathway to respond and adapt to HTTs transmitted from vertebrates to a hematophagous insect responsible for the transmission of Chagas disease.

## Results

### Small RNA profiling in *Rhodnius prolixus*

In order to investigate the dynamics of piRNA expression in oogenesis and early stages of *Rhodnius* development, we generated and sequenced small RNA libraries from embryos (Emb) and 1st instar nymphs (Nym) and analyzed them together with small RNA datasets from previtellogenic stages of oogenesis (PVS) and mature eggs (Egg), that we previously produced [49] (Fig. 1A). After trimming and quality checking, we mapped the reads to the RproC3 version of *Rhodnius* genome available at VectorBase [46,50]. The percentage of reads that mapped to the genome ranged from ∼86% for the PVS1 replicate to ∼95% for the Egg2 replicate (Supplementary Table 1). The length distribution of the reads reveals a bimodal distribution with two major peaks at ∼22 nt and ∼28 nt (Fig. 1B). The former is more prominent in PVS, Egg1 and Nym samples, while the latter is more apparent in Egg2 and Emb samples. To gain insight into the complexity of *Rhodnius* small RNAs, we contrasted our small RNA datasets against *Rhodnius* annotated features available at VectorBase [50]. As expected, a substantial proportion of the mapped reads (i.e. 36.16% for PVS and 65.52% for Nym), match annotated miRNAs (Fig.1C). The abundance of reads corresponding to repetitive sequences shows a reciprocal distribution when compared to the miRNAs. In fact, this category is highly expressed in embryos, while it’s progressively lower in PVS (12.07%) and Nym (10.6%). Once again, the Egg replicates display two non-overlapping distributions with Egg2 being more comparable to the Emb replicates. Other genomic features, including tRNAs, are below 8% in all the samples. A fraction of the reads, ranging from ∼12% in Nym to ∼34% in Emb, could not be classified based on annotated features (i.e. “unassigned”). Next, we profiled the length distribution of the reads associated with transposons and repetitive sequences (Fig. 1D). All the stages analyzed displayed an apparent peak at 21-22 nt, that likely comprises siRNAs, and a ∼28 nt peak corresponding to piRNAs. The abundance of these classes of small RNAs varies in the different samples. The ∼22 nt population is predominant in PVS, while the ∼28 nt is dramatically expressed in embryonic stages and largely exceeds the ∼22 nt set (Fig. 1D). In nymph stages, the levels of ∼22 nt and ∼28 nt small RNAs seem comparable. Once again, the Egg1 and Egg2 replicates provided two different size distributions. Only Egg1 is compatible with maternally loaded piRNAs generated in PVS, while the second replicate seems more similar to the profile obtained in embryos. The easiest explanation for this observation is that the eggs dissected from the adult females used for the Egg2 replicate included a proportion of developing embryos that were retained by the mother. We decided to keep these replicates as they might help shed light on the very initial stages of embryogenesis.

**Figure 1.**
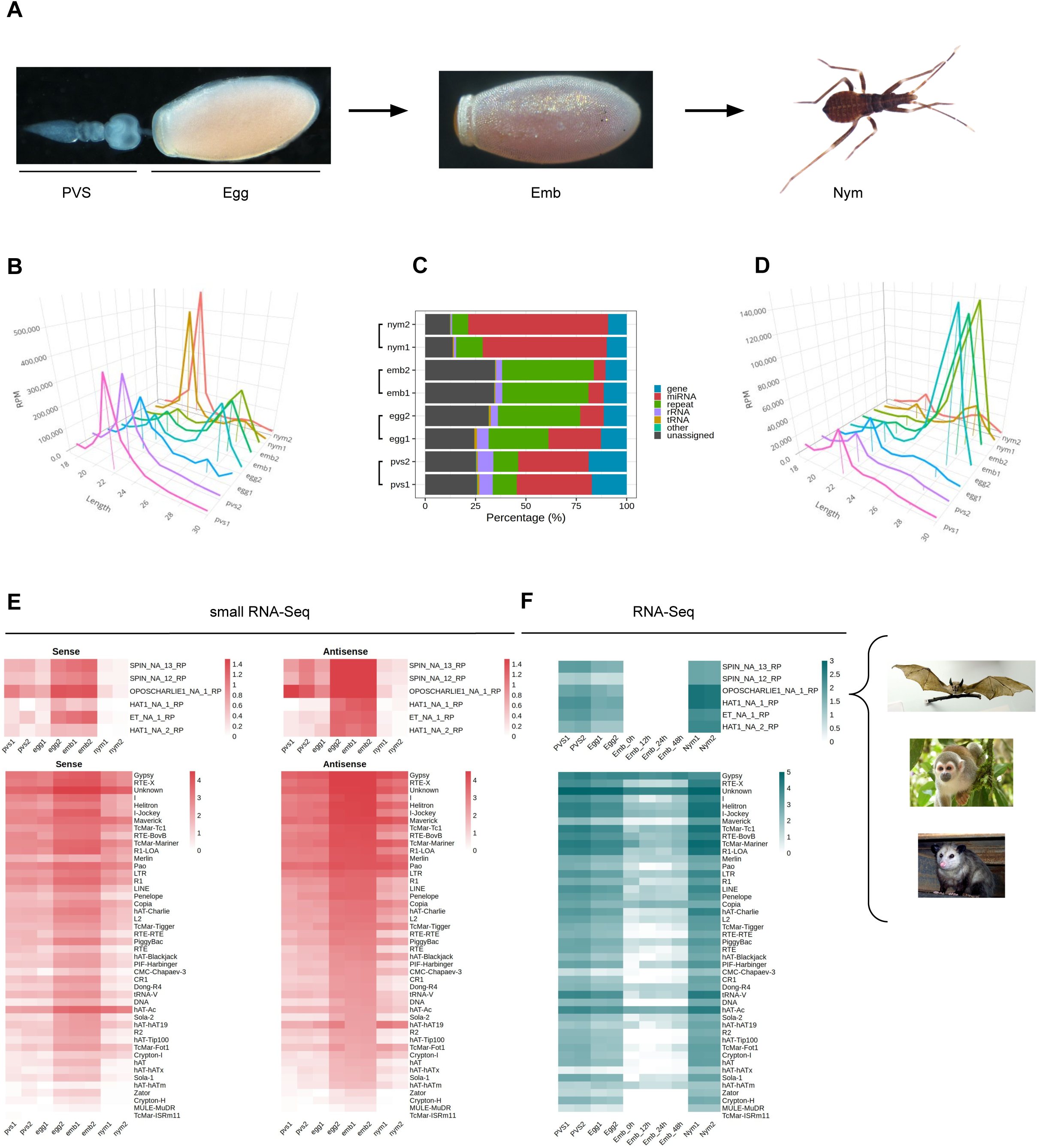
*Rhodnius prolixus* embryos express ∼28 nt long piRNAs targeting Horizontally Transferred Transposons (HTT) and resident TEs. A) Developmental stages of *Rhodnius prolixus* analyzed in the current study. For oogenesis, we investigate the previtellogenic stage (PVS) comprising the tropharium and early follicles independently of the mature chorionated eggs dissected from the female abdomen (Egg). Embryos (Emb) were collected over a 48h period encompassing the cleavage stage until the formation of the head and thoracic segments after gastrulation. 1st instar nymphs (Nym) were also analyzed to investigate postembryonic development. B) Length distribution of all mapped reads for each replicate displayed in RPM. The ∼22 nt and ∼28 nt peaks were anchored to the X-axis (Length) for a better visualization. Y-axis displays RPM values. For each sample, we produced two replicates, which were analyzed separately. B) Small RNA profiling based on percentage of reads mapped to annotated and unannotated (unassigned) regions of *Rhodnius* genome. miRNAs are particularly abundant in oogenesis and postembryonic development (red), while embryonic stages are dominated by repeat-associated small RNAs (green). C) Length distribution of reads mapped to transposable and repetitive sequences of *Rhodnius* displayed in RPM. A major peak comprising ∼28 nt piRNAs is clearly detectable in the two embryo replicates (Emb1 and Emb2) and in one of the mature egg replicates (Egg2). This class of small non-coding RNAs is less abundant in previtellogenic stages of oogenesis (PVS1 and PVS2 replicates), in one of the mature egg replicates (Egg1) and in 1st instar nymphs (Nym1 and Nym2 replicates). D) Heatmaps displaying the expression levels of sense and antisense piRNAs for each TE subfamily in PVS, Egg, Emb and Nym stages. Sense and antisense piRNAs for the horizontally transmitted transposons (HTT) ET, SPIN, OC1 and hAT and for the resident TEs are shown separately. Colorbar values display log_10_(RPM + 1). F) Heatmaps showing TE expression levels as per RNA-Seq in PVS, Egg, Emb and Nym stages. Expression levels for the HTTs ET, SPIN, OC1 and hAT and of the resident TEs are displayed in separate heatmaps. Colorbar values display RPKM. *Rhodnius* acquired the HTTs from opossums, squirrel monkeys and bats (right). For the SPIN and hAT families, Gilbert and coworkers reported two transposon sequences for SPIN and hAT, namely SPIN_NA_13_RP, SPIN_NA_12_RP, hAT_NA_1_RP and hAT_NA_2_RP. One transposon sequence was reported for ET and OC1, which were labeled ET_NA_1_RP and OPSCHARLIE1_NA_1_RP, respectively. For simplicity, we refer to these transposable elements as SPIN, hAT, OC1 and ET throughout the article. Images of bat, squirrel monkey and opossum with Public Domain license were obtained from Wikimedia Commons.

It was shown that four DNA transposon families in *Rhodnius*, that is *SPIN, OC1, hAT* and *ET* originated by horizontal transfer from vertebrate tetrapods [7]. We therefore wondered whether piRNAs related to these HTTs are expressed in *Rhodnius*. To answer this question, we attempted to isolate all the RNAs with 18-30 nt matching HTTs sequences. Strikingly, we find both sense and antisense piRNAs for 6 HTTs reported by Guilbert and collaborators (Fig. 1E) [7]. In addition to the HTTs, 46 transposon families produce both sense and antisense piRNAs with 14 families accounting for more than 90% of the piRNAs. Among them, we could detect abundant piRNAs *LTR/Gypsy*, the LINE elements *Jockey* and *RTE-X*, the rolling-circle *Helitron* and, as expected, for DNA transposons of the *Mariner* family (Fig. 1E). The latter explains 8.9% of the total piRNAs. Among the top most piRNA-producing sequences, we find a class of “Unknown” elements, which might comprise novel transposons or repetitive sequences. In agreement with the general dynamics of the piRNA population in the different stages analyzed, the expression of both sense and antisense piRNAs for the HTTs and other resident TEs appears to dramatically increase during embryogenesis, with the antisense population being more prominent (Fig. 1D and E).

We then analyzed the expression patterns of TEs in oogenesis and early *Rhdonius* development (Fig. 1F). To this aim we combined ovarian and embryonic RNA-Seq datasets that we and others previously generated together with newly sequenced transcriptomes of 1^st^ instar nymphs [48,51]. The temporal expression dynamics of the TEs appear to inversely correlate with piRNA abundance in each stage. The highest TEs expression levels are observed in nymphs, while the lowest are detected in embryonic stages concomitant with peak levels of piRNAs. These results suggest that the 28 nt piRNA population might drive TE silencing during *Rhodnius* embryogenesis, while TE mobilization might be more tolerated in postembryonic development.

### piRNAs in *Rhodnius prolixus* are produced from uni-strand clusters

In order to identify the piRNA source *loci* in *Rhodnius*, we narrowed down our search to piRNAs that map uniquely to the genome [19]. For this study, we adopted a new version of the *Rhodnius* genome generated with the Hi-C technique and available at (https://www.dnazoo.org/assemblies/Rhodnius_prolixus). The Hi-C genome comprises 11 major scaffolds that likely represent the X chromosome and the 10 autosomes of *Rhodnius*. Scaffold 10 was proposed to actually correspond to the X chromosome. We found that unique piRNAs clusterize at 1276 different positions in the genome mainly along the major scaffolds 1-11 (1176) (Fig. 2A). The top 10 clusters account for ∼74% of the total unique piRNAs expressed in *Rhodnius* embryogenesis (Fig. 2B). Only 16 RPCLs (i.e. *Rhodnius prolixus* clusters) display RPM > 100 and 6 of them display RPM > 1000. These RPCL are hosted on scaffold 6 (RPCL1, RPCL3 and RPCL4), scaffold 11 (RPCL2) and the putative X chromosome scaffold 10 (RPCL5 and RPCL6). We find that transposons of the *LINE RTE-X*, *Jockey* and *Gypsy* families together with *Mariner*-like DNA transposons are the most abundant in the top 10 piRNA clusters (Supplementary Fig. 1). It is worth noting however that the most abundant sequence belongs to the “unknown” category and deserves further investigation in the future. The clusters vary in size from a few kilobases to over 700 kb for the RPCL1 locus (Fig. 2C). For the major RPCL1 cluster, we can observe a clear anticorrelation between the levels of piRNAs and cluster transcripts (Fig. 2C). High piRNA levels in embryos coincide with low levels of cluster transcripts. Instead, cluster transcripts appear to accumulate in PVS and, especially, in nymphs, where piRNA expression is at the lowest. Similar expression patterns are observed for the majority of the piRNA clusters (Fig. 2D).

**Figure 2.**
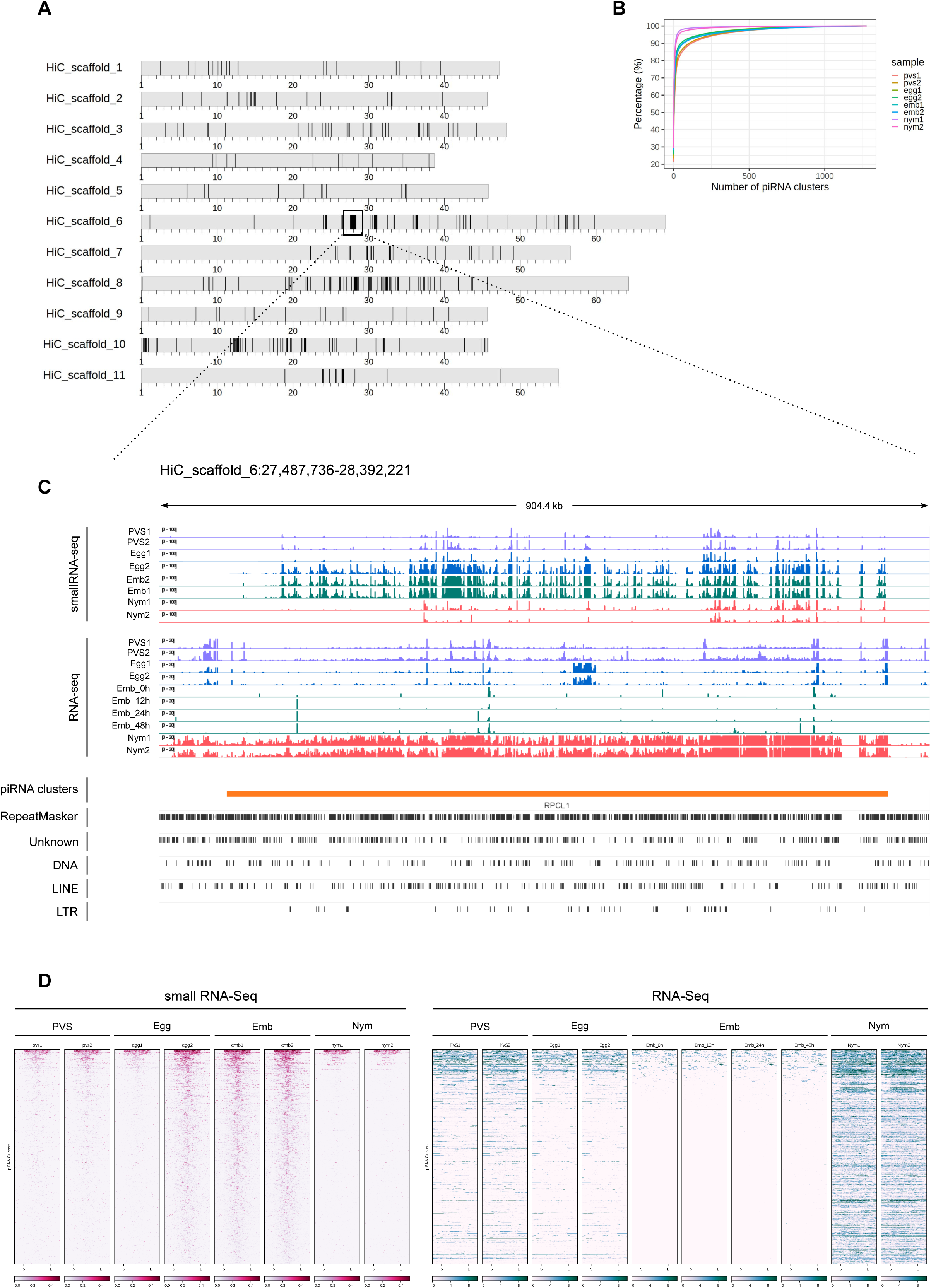
piRNA clusters are a major source of piRNAs during *Rhodnius prolixus* embryonic development. A) Ideograms of the larger scaffolds obtained from the HiC genome assembly that likely represent the 10 autosomes and the X chromosome (scaffold 10) of *Rhodnius*. Each ideogram is scaled to the size of each scaffold, with a scale in megabases. Black vertical lines show the distribution of the piRNA clusters. The major cluster RPCL1 is highlighted (black box).Cumulative percentage of piRNAs in clusters for each replicate. The top 10 RPCLs account for 74% of the total uniquely mapping piRNAs. B) IGV [72] visualization of the RPCL1 cluster region on the HiC_scaffold_6. Uniquely mapped piRNAs for each stage are profiled for each sample (small RNA-Seq profiles). The Y-axis displays RPM (0 to 100 interval). Expression levels of the RPCL1 (RNA-Seq) in all the stages are also displayed. Of note, we included in this analysis the RNA-Seq datasets for embryonic stages 0h, 12h, 24h and 48h post egg laying produced by Pacual and coworkers. The Y-axis displays the RPKM range 0-20. Orange bar highlights the position of the RPCL1 cluster. The distribution of DNA and LTR transposons together with LINE elements and unknown sequences along the RPCL1 region are reported with black boxes. (D) Heatmaps showing the temporal regulation of the cluster-derived mature piRNAs (Small RNA-seq) and piRNA precursor transcripts (RNA-Seq) in PVS, Egg, Emb and Nym stages. Mature piRNA levels inversely correlate with the expression levels of cluster transcripts suggesting that piRNA biogenesis consumes cluster-derived transcripts during embryonic stages. Small RNA-seq values (left side of the panel) are displayed as log_10_(RPM + 1), while RNA-Seq values (right side of the panel) are expressed as RPM. Each line of the heatmap represents a piRNA cluster with the major clusters at the top of the panels.

Next, we investigated the expression of the cluster-derived piRNAs in PVS, Egg, Emb and Nym stages (Fig. 3). Once again, we took into account only uniquely mapping piRNAs. Consistent with the general piRNA temporal profile, the top 10 piRNA clusters display a stark increase in the expression of piRNAs during embryonic stages (Fig. 3A and 3B). Importantly, these *loci* give rise to sense and antisense piRNA populations with the latter generally being more abundant (Fig. 3A and 3B). The alignment of the unique mappers demonstrates that the majority of the clusters are of the uni-strand type (Fig. 3C). Among the major clusters, we were unable to unambiguously identify dual-strand or bidirectional *loci* in *Rhodnius*. RPCL2, which spans over 200 kb, initially appeared to be a bidirectional *locus* in that part of the cluster seems to be expressed from (+) strand and the other from the (-) strand of the genome (Supplementary Fig. 2). Yet, upon closer inspection we realized that the region between the two parts of the cluster with opposite profiles was only partially sequenced. Thus, RPCL2, like other instances of seemingly bidirectional clusters, might represent a genome assembly error, but a more precise characterization will require an improved assembly of the *Rhodnius* genome. We then looked at the orientation of the transposons within the top 20 piRNA clusters (Fig. 3D). Different from the *D. melanogaster flam* locus, where transposon fragments are polarized, *Rhodnius* clusters harbor transposons on both genomic strands. Thus, these *loci* can produce both sense and antisense piRNA, even though transcription occurs from one strand of the genome.

**Figure 3.**
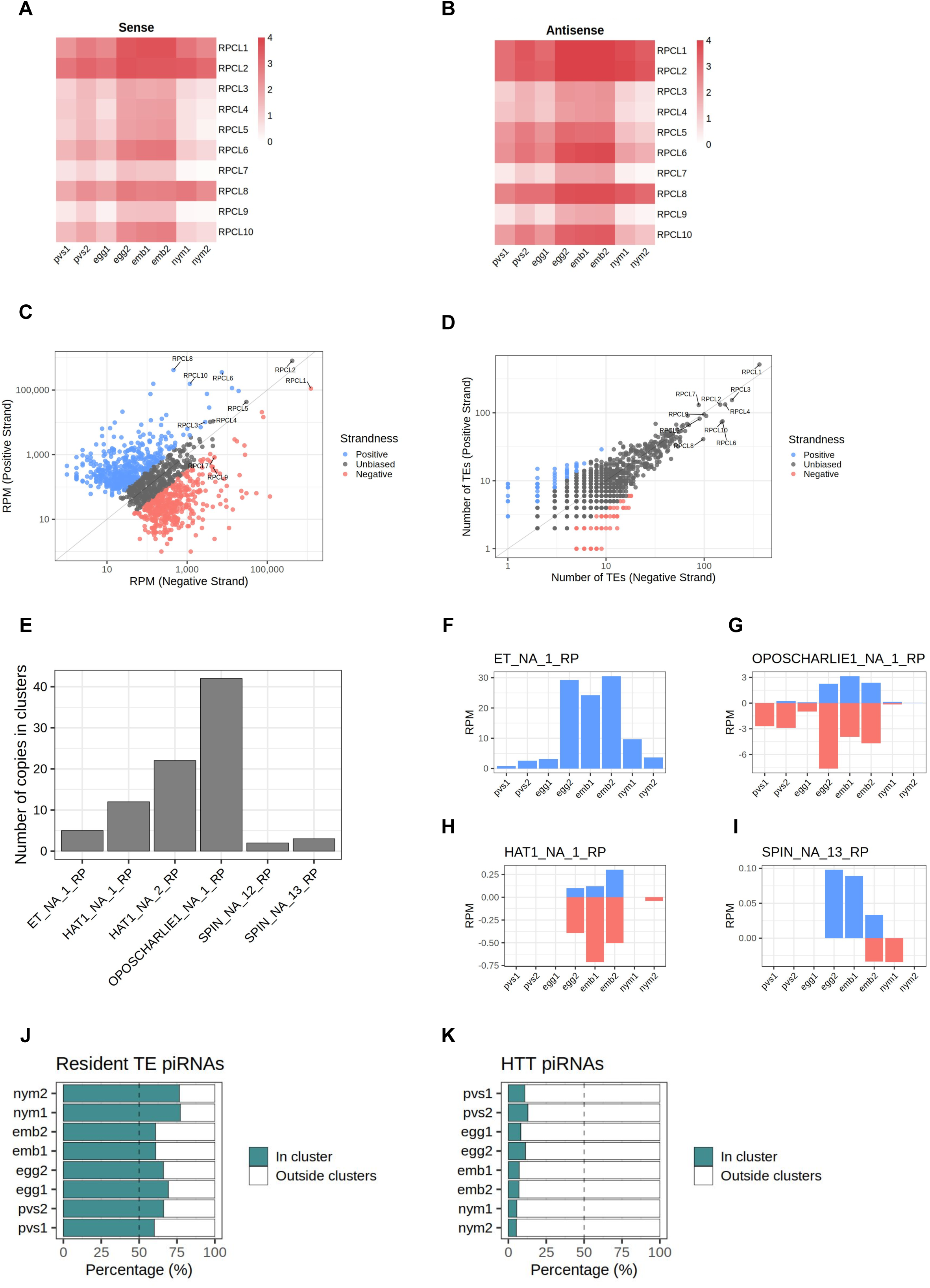
*Rhodnius* piRNA clusters are uni-strand and produce piRNAs against HTTs. A-B) Heatmaps showing sense (A) and antisense (B) piRNA levels for the top 10 piRNA clusters in PVS, Egg, Emb and Nym stages. Colorbar displays log_10_(RPM + 1). C) Scatterplot reporting the strandness of the piRNA clusters. piRNA clusters primarily expressed from the +strand (blue dots) and -strand (red dots) of the genome are displayed. piRNA clusters with apparently unbiased expression are indicated (dark grey dots). A cluster was considered unbiased if less than 75% of its piRNAs could be attributed to a specific strand of the genome. Y-axis values are expressed as RPM. The top 10 piRNA clusters are highlighted. D) Scatterplot showing the orientation of the TE fragment within each piRNA cluster. piRNA clusters harboring TEs predominantly on the +strand (blue dots) or -strand (red dots) of the genome are reported. These categories are mostly formed by small clusters with a low number of TEs. The vast majority of the piRNA clusters are characterized by unbiased distribution of TEs (dark grey dots). The top 10 piRNA clusters are highlighted. E) Number of HTT copies inserted into the piRNA clusters. F-I) piRNAs uniquely mapping to the cluster HTTs. The ET copies (F) seem to exclusively produce sense piRNAs, while OC1 (G), hAT (H) and SPIN (I) generate both sense and antisense piRNAs. Y-axis displays RPM values. X-axis shows the replicates for the PVS, Egg, Emb and Nym samples. J) Percentage of uniquely mapping piRNAs for the resident TEs originating from within or outside the piRNA clusters. K) Percentage of uniquely mapping piRNAs for the HTTs originating from within or outside the piRNA clusters.

Because Rhodnius embryos produce piRNAs targeting the HTTs, we wondered whether these transposons or their fragments are inserted into the piRNA clusters. To answer this question, we aligned the *SPIN*, *hAT*, *OC1* and *ET* sequences with the *Rhodnius* HiC genome and counted the number of occurrences within the piRNA clusters. In agreement with a recent horizontal transfer from vertebrates, the number of HTT copies in the clusters is low and ranges from less than 5 for *ET* and *SPIN* to ∼40 for *OC1* (Fig. 3E). The observation that HTTs sequences are inserted in the major clusters albeit with very low copy numbers prompted us to ask whether these copies are expressed and produce piRNAs. We observe that the *SPIN*, *OC1* and *hAT* sequences generate both sense and antisense piRNAs, while the *ET* element seems to produce mostly sense piRNAs (Fig. 3F-I). These piRNAs however represent only a fraction of the HTT piRNAs. When we compare the cluster-derived piRNAs for resident TEs and HTTs, we observe a striking difference. While for the resident TEs between ∼60% and ∼80% of the piRNAs originate from the clusters (Fig. 3J), less than 15% of the HTT piRNAs are produced from these loci (Fig. 3K). Thus HTT piRNAs are mostly generated by the transposon copies scattered in the genome.

### HTT piRNAs display unique features

In order to investigate the characteristics of the HTT piRNAs, we first profiled the piRNAs along the transposon sequences. Their distribution is not homogenous and they seem to accumulate at certain positions (Fig. 4A-D). We then investigated the overlap between the 5’ ends of sense and antisense sequences for the paired piRNAs (Fig. 4A’-D’). The 10 nt overlap typical of the ping-pong mechanism is either modest or it is confused among other peaks, as in the cases of *OC1* and *ET* (Fig. 4A’ and 4B’). Instead, the *SPIN* and *hAT* piRNA mates display an apparent ∼20 nt overlap (Fig. 4C’ and 4D’), that is characteristic of siRNA heteroduplexes produced by the activity of the Dcr2 enzyme. Yet, these piRNA pairs are over 28 nt long. We next investigated the nucleotide distribution frequencies along the HTT piRNAs. We find that both sense and antisense piRNAs display a 1U bias, while the 10A bias in sense piRNAs is less apparent (Fig. 4A“-D”). The most abundant piRNA pairs for *hAT* and *Spin* further underscore the unique properties of the HTTs piRNAs (Fig. 4C-D inset). These pairs comprise piRNAs of over 28 nt in length, which overlap by 19 nt and 21 nt respectively. Remarkably both sense and antisense piRNAs display 1U and 10A residues for the *hAT* pair, while only the sense *Spin* piRNA has a 10A residue.

**Figure 4.**
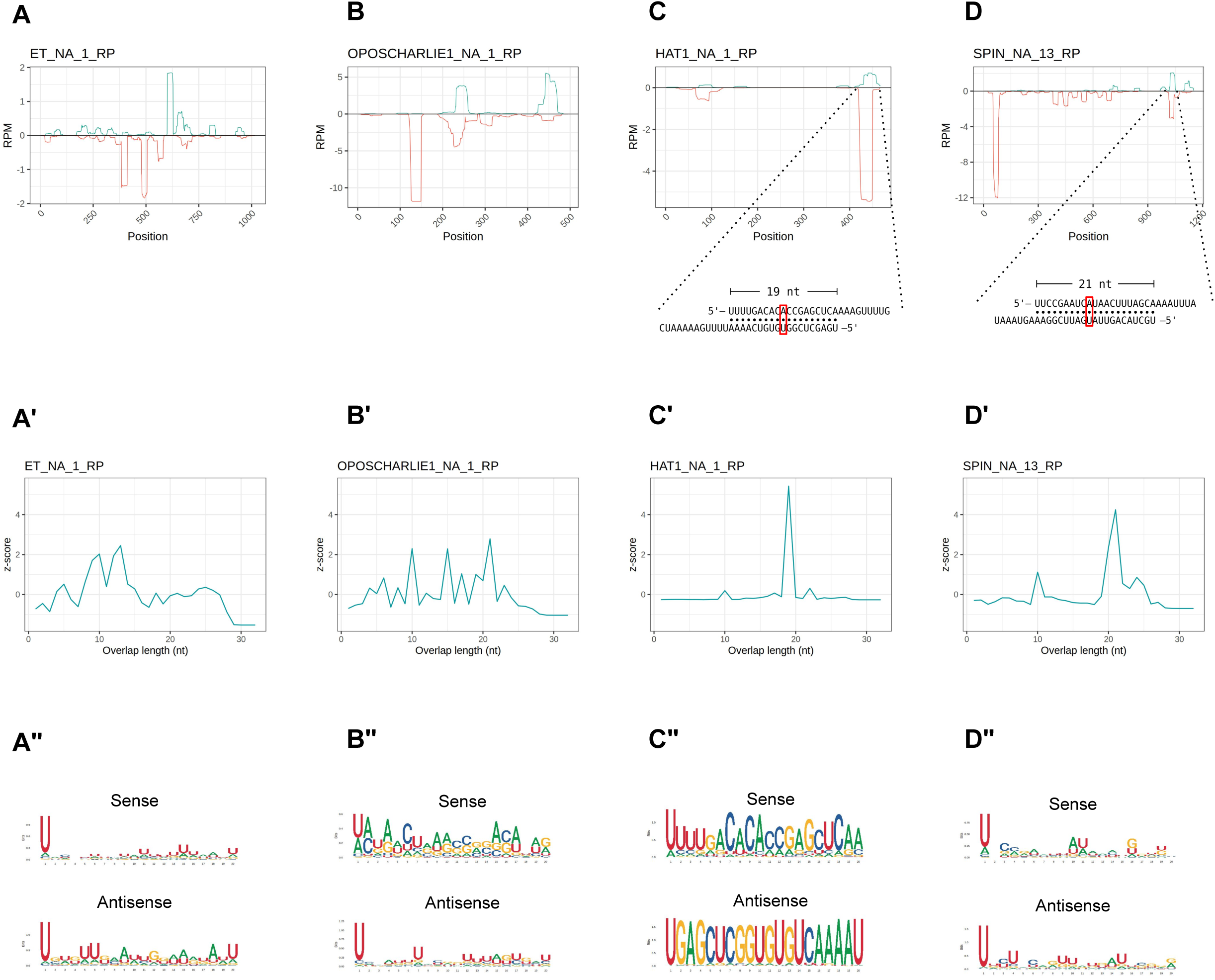
HTT piRNAs do not engage in a canonical ping-pong amplification mechanism. A-D) piRNA profiling along the sequences of each HTT with sense piRNAs displayed with positive RPM values (green) and antisense piRNAs with negative RPM values (red). For the hAT and SPIN elements, the most abundant sense/antisense piRNA pairs are shown. Of note, the sense piRNAs pairs do not display the typical ping-pong signature, but overlap by ∼20 nt. Yet, the sense mate for each pair bears an Adenine residue at the 10th position (red box). A’-D’) 5’ to 5’ overlap length frequency reported as z-score values for each HTT. A“-D”) Sequence logos displaying nucleotide bias for sense and antisense piRNAs mapped to the HTTs.

### piRNAs in *Rhodnius prolixus* display ping-pong and phasing signatures

The HTTs have been suggested to have invaded the *Rhodnius* genome over the past 50 million years. Thus, the piRNAs associated with these elements likely display peculiar characteristics that are not shared by other resident TEs. To shed light on this aspect, we filtered all the small RNA sequences ranging from 24 to 30 nt in length, which yielded ∼36 million putative piRNA sequences (Supplementary Table 2). We then investigated whether a ping-pong amplification system is active during *Rhodnius* development. (Fig. 5). First, we estimated the fraction of paired piRNAs in all the stages and found that it ranged from ∼19% in PVS to ∼29% in Embryos (Fig. 5A). As expected the proportion of unpaired piRNAs is more abundant and likely comprises primary piRNAs. Different from the HTTs, we found that a substantial fraction of the pairs (∼37%) overlap by exactly 10 nt at their 5’ ends (Fig. 5B). The analysis of nucleotide distribution shows that antisense piRNAs display a clear 1U bias in all the samples and that sense piRNAs are enriched with a 10A residue (Fig. 5C). Sense piRNAs also bear a 1U bias, which was also described in *Tribolium castaneum*, although its meaning remains unclear. Thus, piRNAs engage in a ping-pong amplification mechanism in *Rhodnius,* which is not restricted to the germline as in *D. melanogaster*, but is also active in embryos and 1^st^ instar nymphs. Previous studies demonstrated that piRNA production via ping-pong cycle is coupled with production of phased primary piRNAs in a range of different animals [30,31]. To address whether primary piRNAs can be produced in a phased manner, we compared the distances between 3’ and 5’ ends of different piRNAs mapped to the genome (Fig. 5D). We find that a high frequency of consecutive piRNAs (Z_0_ > 9) can be detected in all the stages . Therefore, phased primary piRNAs are produced in *Rhodnius prolixus* and likely share a similar mechanism of biogenesis as observed in other animals.

**Figure 5.**
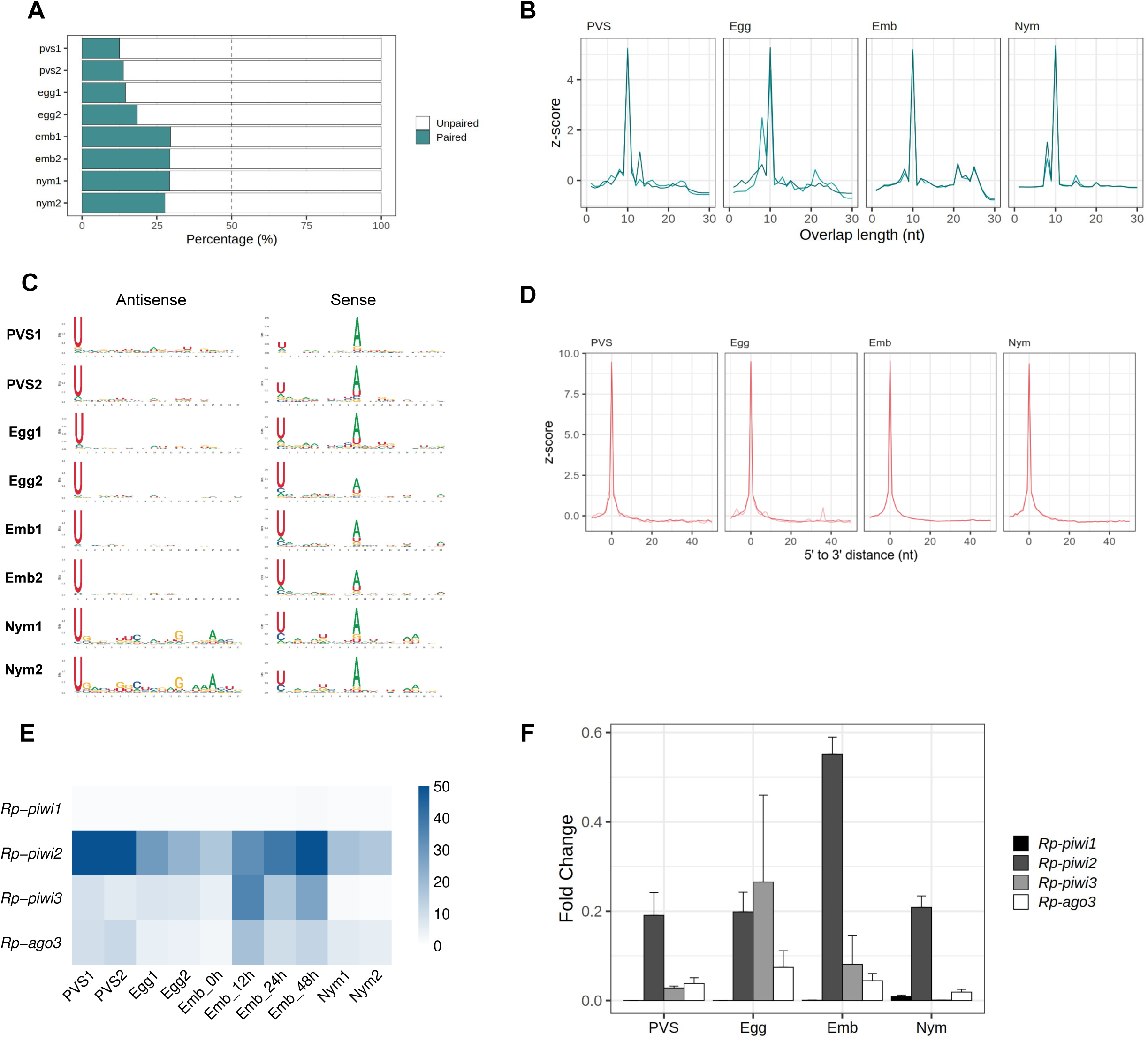
The ping-pong amplification loop is active in *Rhodnius prolixus*. A) Percentage of paired and unpaired piRNAs for PVS, Egg, Emb and Nym replicates. B) 5’ to 5’ overlap length frequency displayed in z-score for piRNAs mapped to resident transposons for each stage. C) Sequence logo showing nucleotide bias for each position of antisense and sense piRNAs mapped to transposable elements in all the replicates. D) Phasing quantification in z-score showing the 5’ to 3’ distance between consecutive piRNAs for each stage. E) RNA-Seq analysis of the *Rp-PIWI* gene expression in oogenesis and early *Rhodnius* development. Colorcode displays RPKM values. F) qRT-PCR analysis of the temporal expression levels of the *Rp-PIWI* genes during oogenesis and early stages of *Rhodnius prolixus* development. PVS=previtellogenic stage, Egg=mature chorionated eggs, Emb=embryos and Nym=1st instar nymphs. Y-axis shows the mean Fold Change. Error bars display standard deviation from the mean.

The discovery of a ∼28 nt piRNA population in embryos and their involvement in a ping-pong amplification loop prompted us to investigate what *Rp-PIWI* genes are expressed in this stage as well as in first instar nymphs. We have previously shown that *Rp-piwi2*, *Rp-piwi3* and *Rp-ago3*, but not *Rp-piwi1* are expressed in and important for oogenesis in *Rhodnius* [47]. In order to expand this analysis, we combined *in silico* and wet biology analyses (Fig. 5E and F). First, we interrogated our RNA-Seq datasets together with those produced by Pascual and Rivera-Pomar [51]. Consistent with our previous studies, *Rp-piwi2*, *Rp-piwi3* and *Rp-ago3* are expressed in *Rhodnius* oogenesis (Fig. 4E) [47]. Interestingly, these genes are also upregulated in the first 48h of embryonic development, while their levels drop substantially once the 1^st^ instar nymphs hatch (Fig. 4E). During postembryonic development, the levels of *Rp-piwi3* are below detectable levels. *Rp-piwi1* is apparently not expressed in any of the stages analyzed. In order to validate these observations, we performed RT-qPCR assays on total RNA extracted from PVS, Egg, Emb (0-48h) and Nym stages. We used oligonucleotide pairs specific for each *Rp-PIWI* gene and for *Rp-rp49*, which served as internal control (Fig. 5F) [47]. The Fold Changes of each gene are in line with RPKM values. Interestingly, the *Rp-piwi2* levels dramatically increase in embryogenesis compared to the other stages analyzed and largely exceed those of the remaining *Rp-PIWIs* pointing to an important role in the biogenesis and/or function of the ∼28nt piRNA population. Of note, the expression of *Rp-piwi1* as per RT-qPCR is low, but clearly detectable in 1^st^ instar nymphs (Fig. 5F). Whether *Rp-piwi1* transcripts exert any functions in postembryonic development remains to be elucidated. Our results reveal that *Rp-PIWI* genes are therefore temporally regulated during oogenesis and early *Rhodnius* development.

### TE siRNAs are predominantly expressed in oogenesis

Although the piRNA pathway represents a major surveillance system keeping mobile elements in check, it was shown that Dcr2-dependent siRNAs control transposon silencing in *Drosophila* somatic cells [52]. To investigate whether this mechanism is conserved in *Rhodnius*, we interrogated the transposon-related sequences in the 18 to 23 nt size range and analyzed the ping-pong signal, nucleotide bias and strandness (Fig. 6). We find a clear population of sense and antisense small RNAs centered on ∼22 nt in length that is particularly abundant in PVS and Egg compared to the remaining samples (Fig. 6A and B). Surprisingly, the two Egg replicates display remarkably similar siRNA profiles. Together with the piRNA analysis, this observation suggests that the Egg2 replicate might be capturing the early embryonic stages where the siRNA levels are still high, but the piRNA production from the zygotic genome is building up. The degree of overlap between sense and antisense sequences reveals that the ∼22 nt small RNAs expressed in PVS, Egg and to a lesser extent in Emb and Nym are siRNAs, while Emb samples comprise ∼22 nt piRNAs (Fig. 6C). Consistent with this, the nucleotide distribution shows that the 1U together with a modest 10A bias are detected only in embryos, but not in the other stages analyzed (Fig. 6D). Finally, we wondered whether TE siRNAs can originate from the piRNA clusters. The analysis of uniquely mapping reads reveals that a substantial proportion of siRNAs can indeed be produced by these *loci* in all the stages (Fig. 6E). Accurate quantification of siRNAs produced from clusters versus TEs or other regions of the genome is confounded by the presence of piRNAs in the same size range. Surprisingly, we do not find sequences corresponding to the HTTs in the siRNA population. These observations reveal that siRNAs and piRNAs expression patterns are temporally regulated and partially overlap during oogenesis and early stages of *Rhodnius* development.

**Figure 6.**
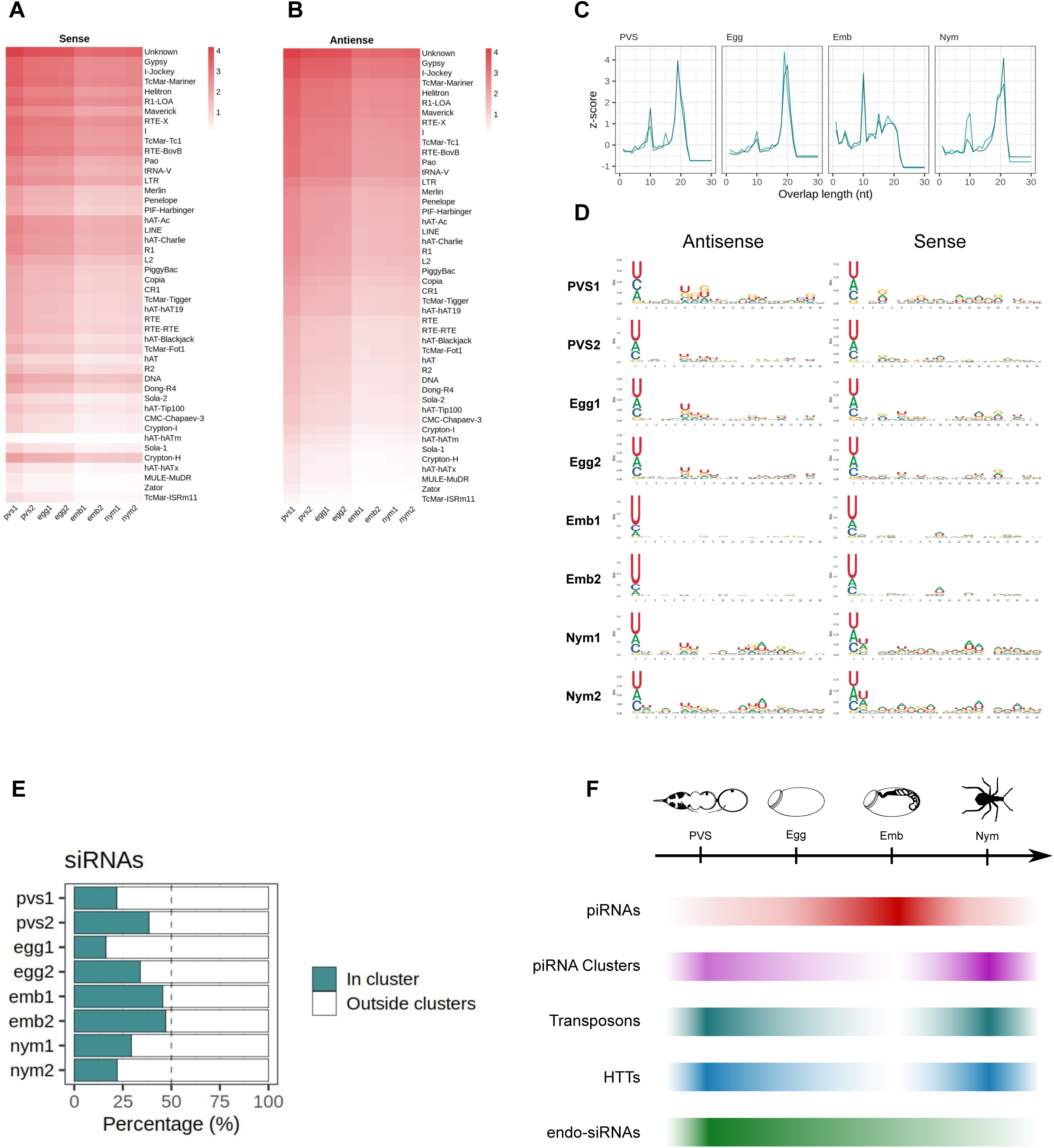
piRNA clusters generate ∼22nt siRNAs in *Rhodnius prolixus*. A-B) Heatmaps showing sense (A) and antisense (B) ∼22 nt long siRNAs mapped to the *Rhodnius* TE subfamilies. Colorbar displays log_10_(RPM + 1). siRNAs appear to be expressed at higher levels in oogenesis compared to embryonic and postembryonic stages. C) Z-scores of the 5’ to 5’ overlap length frequency for PVS, Egg, Emb and Nym samples. (D) Sequence logos showing nucleotide bias for each position of antisense and sense siRNAs for each stage. The 1U bias typical of the piRNAs is only detectable in embryos. E) Percentage of uniquely mapping siRNAs originating from the piRNA clusters (green bars) versus other regions of the genome (white) for PVS, Egg, Emb and Nym stages. F) Schematic diagram depicting the dynamics of piRNAs (red bar), piRNA cluster transcripts (purple bar), TEs (dark green), HTTs (blue bar) and siRNAs (light green bar) during oogenesis, embryogenesis and postembryonic development of *Rhodnius prolixus*.

## Discussion

Studies in *Drosophila melanogaster* and its sibling species highlighted the adaptability of the piRNA pathway to newly invading transposons [53]. Aside from the Drosophilids however, how animal cells respond to TE invasion is still largely unknown. In this study, we show a striking example of piRNA pathway adaptation to TEs that were transferred from vertebrate tetrapods to the assassin bug *Rhodnius prolixus*. *Rhodnius* belongs to the Triatomine subfamily and has been firmly connected to the transmission of Chagas disease. It was shown that the *SPIN*, *OC1*, *hAT* and *ET* elements were horizontally transmitted to *Rhodnius* from bats, opossums and squirrel monkeys [7]. Because of the hematophagous habit of *Rhodnius* however, HTT might have occurred while the insect was feeding on the blood of the vertebrate hosts. This event provided the unique opportunity to investigate the response of the piRNA pathway to transposons exchanged between two different phyla in the past 50 million years. When we compare the complement of piRNAs targeting resident transposons and HTTs in *Rhodnius*, we find some similarities, but also puzzling differences. A major population of ∼28 nt long piRNAs is expressed at high levels during *Rhodnius* embryogenesis and comprises sequences of 46 transposon families. Some examples are the *Gypsy*, *RTE-X*, *Helitron*, *Maverick*, *I*, *Penelope* and the abundant class of *Mariner* transposons [8]. We have referred to all these transposons as resident TEs, but it should be noted that they are frequently involved in horizontal transfer events. For these elements, we find primary piRNAs with the typical 1U bias and secondary piRNAs with Adenine at 10th position (10A) for the sense piRNAs. Also, consistent with an active ping-pong amplification mechanism, sense and antisense piRNA mates overlap by 10 nt. Similar to more basal holometabolous and hemimetabolous insects, but unlike *D. melanogaster*, the ping-pong signal is not restricted to the gonads in *Rhodnius*, but it is especially active in embryonic and in post-embryonic developmental stages. The piRNA complement targeting the HTTs instead displays unique properties. Even though HTT piRNAs are ∼28 nt long and display a 1U bias, the ping-pong mechanism is ambiguous. The 10 nt overlap is modest or confused among other signals. Instead, the most abundant *SPIN* and *hAT* piRNA mates display a ∼20 nt overlap at their 5’ ends. Yet, the 10^th^ position of the sense piRNAs is an Adenine as observed for canonical ping-pong piRNA pairs. These observations suggest that the biogenesis of HTT piRNAs in *Rhodnius* differs from the more canonical mechanism that drives resident TE piRNA maturation and that was described in detail in the fruit fly *D. melanogaster*. The lack of ping-pong signature for the HTTs might reside at least in part in the genomic sources of their piRNAs. In this study, we show that piRNA clusters in *Rhodnius* are mostly of the uni-strand type, but unlike the *Drosophila flam* locus [19], their TEs are not polarized. For this reason, *Rhodnius* clusters can produce both sense and antisense piRNAs, even though the transcription occurs from one genomic strand. We find that more than 50% of the unique piRNAs targeting resident TEs originate from the piRNA clusters, while these *loci* account for less than 15% of the HTT piRNAs. This observation is consistent with the low number of HTT copies inserted into the clusters. These sequences therefore, might not produce sufficient levels of antisense transcripts to robustly activate the ping-pong mechanism. The *ET* element provides an extreme case, whereby all the cluster copies generate sense piRNAs, thus preventing the production of secondary piRNAs. How the *hAT* and *SPIN* piRNA mates overlapping by ∼20 nt are generated and what factors are involved in the mechanism remains to be elucidated. Interestingly, our results reveal that *Rhodnius* expresses siRNAs for resident TEs, but not for the HTTs. Hence, primary piRNAs produced from HTT copies or fragments scattered in the genome, rather than siRNAs, might provide the first line of defense against newly invading transposons in this insect species.

TE siRNA are expressed from piRNA clusters and TE copies dispersed in the genome during oogenesis and to a lesser extent during embryonic and postembryonic stages. RNAi might contribute to reducing transposon transcripts during oogenesis, although it is unable to completely silence these elements [48]. It was recently shown that maternally provided siRNAs loaded in the developing eggs promote piRNA cluster activation in the *Drosophila* embryo [54]. Unlike *Drosophila*, *Rhodnius* does not produce abundant piRNAs during oogenesis. Thus in *Rhodnius* and other more basal insects, the siRNA complement might be especially critical for jumpstarting piRNA production in early embryonic stages. Transposon downregulation is subsequently taken over by the ∼28 nt piRNA pool during embryogenesis. Transposon transcripts maternally deposited in the developing oocytes might feed the ping-pong amplification loop at this stage together with cluster-derived antisense piRNAs. piRNA biogenesis seems to result in the consumption of the cluster-derived piRNA precursors and the downregulation of TEs. In post-embryonic stages instead both siRNAs and piRNAs are reduced, while TE and piRNA cluster transcripts appear to accumulate, thus pointing to a minor sensitivity of the 1^st^ instar nymphs to transposon expression. In support of these conclusions, the levels of piRNAs and siRNAs seem to inversely correlate with those of TEs and piRNA cluster transcripts in all the stages analyzed (Fig. 6F).

*Rhodnius* harbors four *PIWI* genes and the knockdown of *Rp-piwi2*, *Rp-piwi3* and *Rp-ago3* by parental RNAi results in partial or complete female adult sterility [47,55]. Our assays show that these *Rp-PIWI* genes are expressed in embryos and nymphs with *Rp-piwi2* being dramatically upregulated in embryos coinciding with peak levels of ∼28 nt piRNAs. Thus, *Rp-piwi2* likely exerts a crucial role in piRNA biogenesis or function. Since Ago3 was shown to coordinate the ping-pong cycle together with Aub in *Drosophila* [18,19], it is tempting to speculate that the protein products of the only *ago3* ortholog in *Rhodnius* (i.e. *Rp-ago3*) and *Rp-piwi2* genes might regulate a similar mechanism in *Rhodnius* embryos. However, *Rp-ago3* might actually act in concert with *Rp-piwi3* in this mechanism. The observation that *Rp-piwi3* expression levels are low in oogenesis and almost undetectable in nymphs, where the ∼28 nt piRNAs are barely produced, seem to lend support to this hypothesis. The cell lines that were recently developed from *Rhodnius* embryos will likely provide a valuable tool to dissect the function of the different *Rp-PIWI* genes in piRNA biogenesis and transposon silencing as well as shed light on the transcriptional regulation of the piRNA clusters [56].

### Conclusions

Horizontal transposon transfer has been well characterized in Drosophilids, where numerous instances of P-element transfer from *D. willinstoni* and *D. simulans* to *D. melanogaster* were described [57,58]. Fruit flies therefore provided the opportunity to dissect the response of the piRNA pathway to P-element invasion [59]. To our knowledge however, this is the first study to link piRNAs to HTTs exchanged between vertebrates and an hematophagous insect of medical interest. Studies in *Rhodnius* will therefore help shed light on the evolution of the piRNA pathway and on the initial steps of the arms race between invading TEs and small RNA-based defense systems.

## Methods

### Ethic statement and insect handling

All the experimental protocols involving animals were carried on in accordance with the guidelines of the Committee for Evaluation of Animal Use for Research (Universidade Federal do Rio de Janeiro, CAUAP-UFRJ) and the NIH Guide for the Care and Use of Laboratory Animals (ISBN 0-309-05377-3). Protocols were approved by CAUAP-UFRJ under registry #IBQM155/13. Specialized technicians at the animal facility (Instituto de Bioquímica Médica Leopoldo de Meis, UFRJ) handled the rabbit husbandry under strict guidelines and the supervision of veterinarians.

Insects were maintained at controlled temperature (28°C) and relative humidity (70-80%) and fed with rabbit blood at regular intervals to promote ecdysis and oogenesis. After the first blood feeding, all adult insects were fed every 21 days. Dissection of the adult females, where necessary, was carried on 7 days after the first or second blood meal. All animal care and experimental protocols were performed according to the ethics guidelines.

### Total RNA extraction and small RNA purification

Total RNA for small RNA library preparation was extracted from 30 embryos and 20 1^st^-instar nymphs for each biological replicate. Embryos were collected over a two-day period spanning the initial stages of embryogenesis up to the formation of head and thoracic segments [60]. Two biological replicates were produced for each developmental stage. Ovaries were dissected in ice-cold 1X PBS and immediately stored on ice. Previtellogenic tissues and mature eggs were teased apart under light microscopy and subjected to total RNA extraction using Trizol Reagent (Invitrogen) as per manufacturer’s instructions. For the production of small RNA libraries, 20 µg of total RNA was separated on polyacrylamide gel containing 15M Urea (Pane et al., 2011). A gel slice corresponding to 18 to 30 nucleotide RNAs was manually retrieved from the gel and the RNAs were eluted in 3M NaCl overnight. Finally, RNA was precipitated using 0.7 volumes of Isopropanol and 1 ul Glycogen and resuspended in RNAse-free water. Small RNA library preparation and sequencing were carried out at the Max Planck Institute facility in Freiburg (Germany) with Truseq Small RNA library preparation kit as per manufacturer’s protocol (Illumina). Total RNA extracted from 1^st^ instar nymphs was also used to produce and sequence RN-Seq libraries (Macrogen, Korea). RNA-Seq datasets corresponding to four embryonic stages of *Rhodnius* development were produced by others and are available at NCBI-SRA [51] (SRR14606474, SRR14606475 and SRR14606473).

### Bioinformatic analysis

Small RNA-Seq libraries were subjected to adaptor removal and trimming using TrimGalore! [61]. Reads with length lower than 18 nt were filtered out. Trimmed reads were mapped with Bowtie v1.2 [62] to *Rhodnius prolixus* reference genome available at VectorBase [50] allowing up to 3 mismatches and no more than 50 mappings per read (-v 3 -m 50 -a --best --strata). Qualities were assessed with FastQC software [63], genomic coverages were compared with deepTools [64] and final quality reports were generated using multiQC [65]. Library characterization was performed using featureCounts from the subread package [66], where counts to each annotated feature were normalized by number of mappings and overlapping features (-O -M --fraction). Repeats and TEs sequences were de novo identified using RepeatModeler v2.0.1 [67]and genomic regions obtained with RepeatMasker v4.1.2 [68]. Sequences of the HTTs have been previously published [7]. Reads overlapping with TEs, with length distribution of 24 to 32 nucleotides and not overlapping with miRNAs, rRNAs, snRNAs, simple repeats or low complexity regions were characterized as piRNAs. Sense and antisense piRNAs were separated by overlapping piRNA mappings with TEs annotation using bedtools intersect and forcing strandness (-s and -S). Fasta sequences were extracted using samtools [69] and sequence logos constructed using ggseqlogo [70]. For ping-pong signal analysis, each sense piRNA was used to search for 5’ to 5’ overlapping antisense piRNAs and the size of the overlap counted for each match. Reads mapped to multiple places in the genome were apportioned. Phasing analysis was done as previously described [30]. For piRNA cluster identification, the chromosome-length genome assembly of *Rhodnius* (https://www.dnazoo.org/assemblies/Rhodnius_prolixus) was used and all the steps described previously were applied to this version as well. First, annotations of a sliding window with 5 kb in size and steps of 1 kb were generated using the makewindow tool from bedtools v2.31 [71]. Then, bedtools coverage was used to access the uniquely mapped piRNA coverage in each window and only windows with a density greater than 5 piRNAs/kb covering at least 12% of the window were retained. Finally, overlapping windows were merged and merged windows with a distance less than 20 kb from each other were combined to produce the final piRNA clusters. RNA-seq data were pre-processed and aligned using snakePipes 62. Genes and transposable element read counts were produced using featureCounts allowing fraction counts for overlapping features. Final expression was calculated as Reads Per Kilobase per Million (RPKM).

### Quantitative Rt-PCR assays

For quantitative Rt-PCR assays, 1 ug of total RNA extracted with Trizol (Invitrogen) was submitted to DNAse treatment with Turbo DNAse free kit (Ambion) and cDNA synthesis using the MultiScribe Reverse Transcriptase (Thermo Fisher Scientific) as per manufacturer’s instructions. The resulting cDNAs were used for qPCR assays with the set of oligonucleotides previously described [47]. Roughly 50 ng of cDNA for each sample was mixed with oligonucleotides specific to target genes and SYBR green reagent (Life Technologies). RT-qPCR was carried on QuantStudio 7 Flex (ThermoFisher). Fold change differences were plotted as 2^-ΔCt^.

## Acknowledgments

We would like to thank Bernardo Carvalho, Pedro Lagerblad de Oliveira, Ana Cristina Bahia Nascimento, Marcos Farina de Souza e Marcia Cury El-Cheikh for their constant support. We are grateful to Eric Aguiar and members of the Functional Genomics Laboratory for helpful suggestions and critical reading of the manuscript. We thank Kelli Cristina Melquiades Mendes, Daniela Sodré Leal and Graciela Venturi for the invaluable technical support.

## Data availability

Small RNA sequencing datasets from embryonic (Emb1 and Emb2 replicates) and 1st instar nymphs (Nym1 and Nym2 replicates) were deposited at NCBI SRA database (accession numbers ).

## Author contributions

Tarcísio Fontenele de Brito: Data curation, Formal analysis, Investigation, Methodology, Validation, Visualization, Writing – original draft, Writing – review & editing. Maira Arruda Cardoso: Data curation, Investigation, Methodology, Validation, Writing – original draft, Writing – review & editing. Nazerke Atinbayeva: Investigation, Writing – review & editing. Ingrid Alexandre de Abreu Brito: Investigation, Methodology, Writing – original draft. Lucas Amaro da Costa: Methodology, Investigation, Writing – original draft. Nicola Iovino: Data curation, Funding acquisition, Resources, Writing – original draft, Writing – review & editing. Attilio Pane, Conceptualization, Data curation, Formal analysis, Funding acquisition, Investigation, Methodology, Project administration, Resources, Supervision, Validation, Visualization, Writing – original draft, Writing – review & editing.

---

**Supplementary Fig 1. Abundance of transposon families in the top 10 piRNA clusters.** X-axis displays the number of elements found for each transposon family (Y-axis) in each of the top 10 piRNA clusters. Since the “Unknown” elements are the most abundant ones, they are shown separately at the bottom.

**Supplementary Fig 2. Partially sequenced region in RPCL2.** Positive (blue) and negative (red) read coverage along the piRNA cluster RPCL2 (top). At first inspection, RPCL2 seems like a bidirectional cluster. However, when looking at the exact region where the expression shifts (bottom), a partially sequenced region is notable and might represent an assembly error.

**Supplementary Table 1. General mapping statistics**

**Supplementary Table 2. piRNA identification and pairing information**

