## Supplementary Material for "“Embryonic piRNAs target horizontally transferred vertebrate transposons in assassin bugs”"

### Supplementary Figure 1

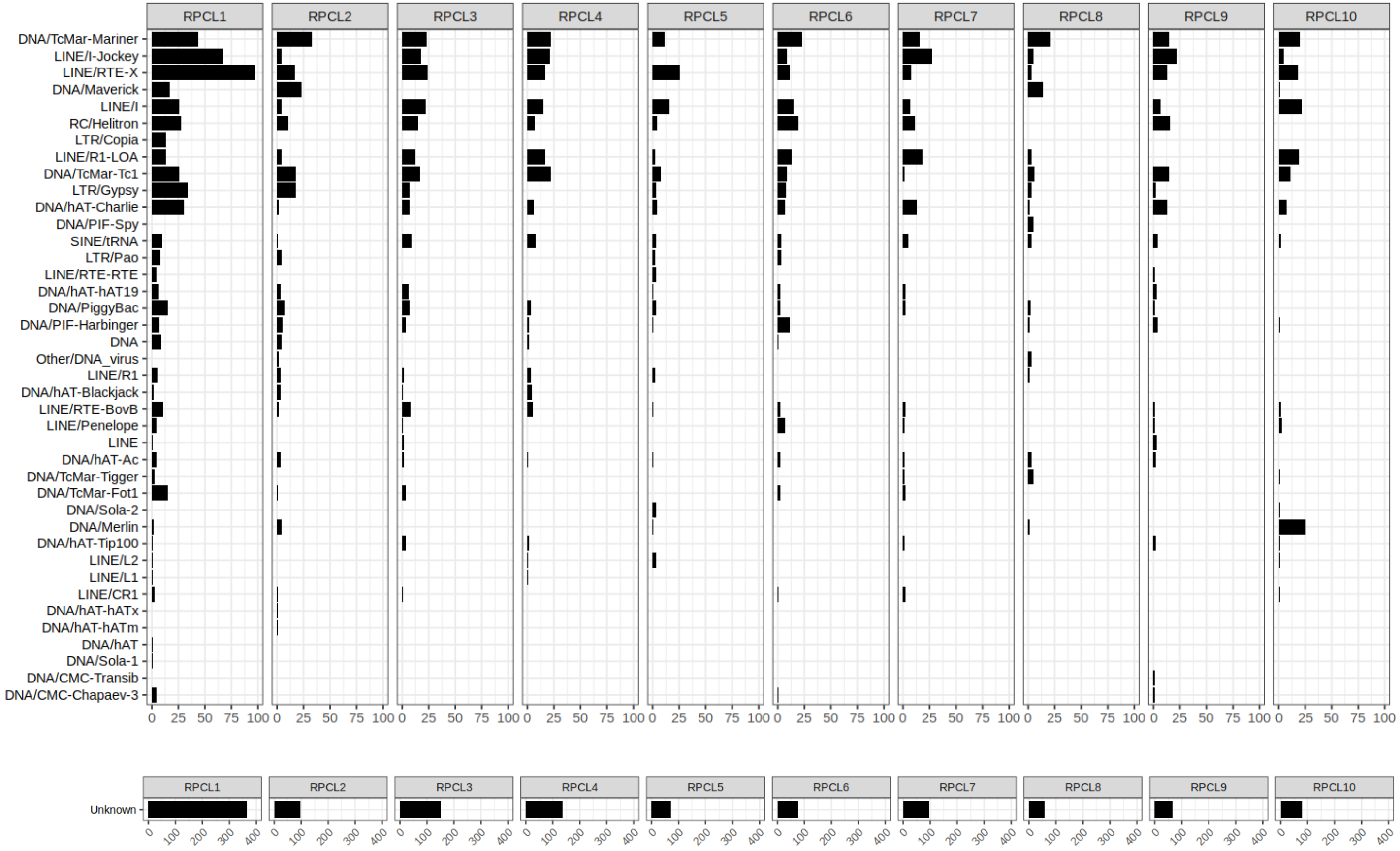

**Supplementary Figure 2**

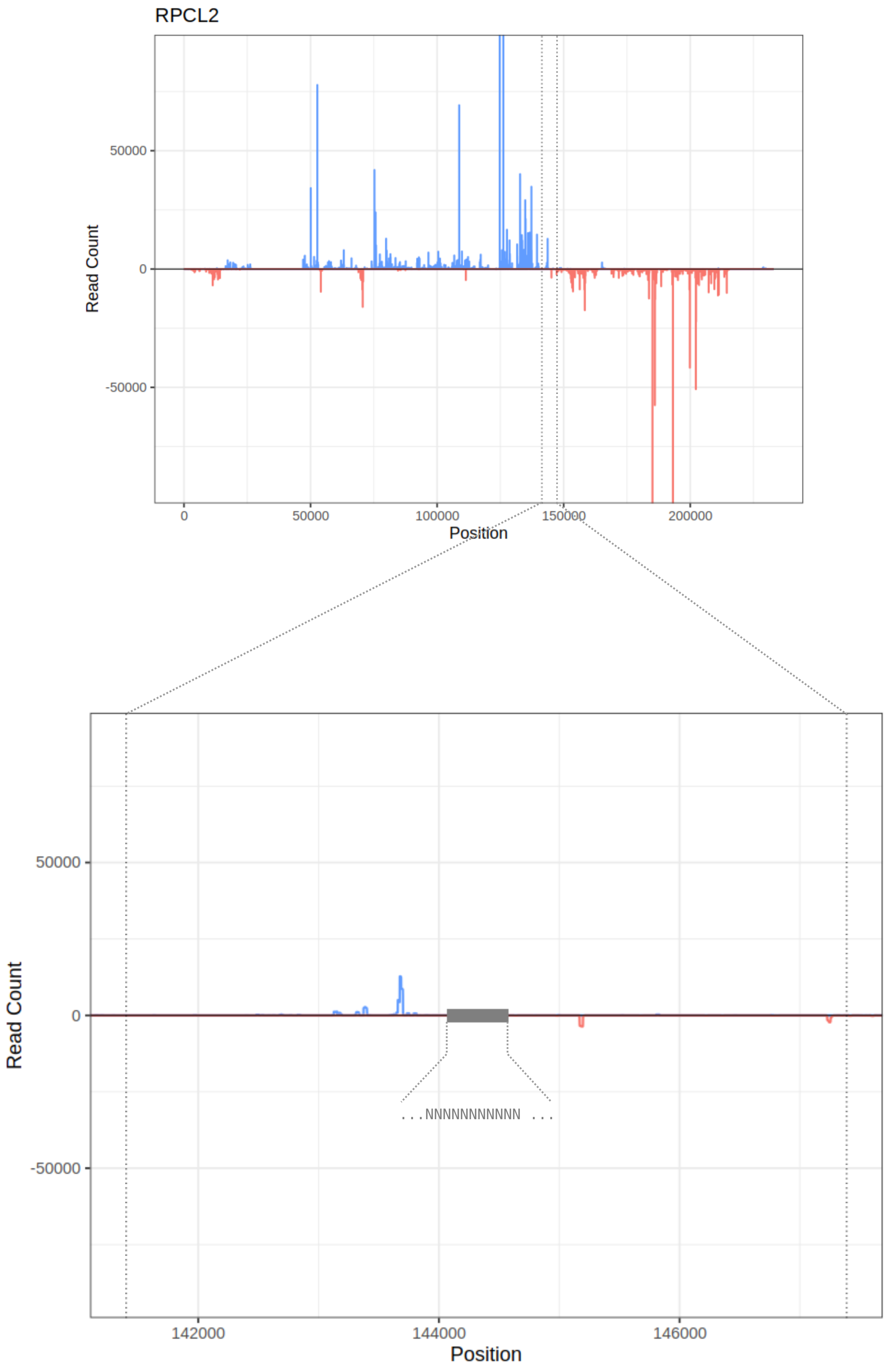

**Supplementary Table 1**

| Sample | Total Reads | Total Reads After QC | | Reads mapped to RproC3 | | Reads mapped to HiC assembly | |
|---|---|---|---|---|---|---|---|
| PVS1 | 5.582.939 | 4.277.071 | 76,61% | 3.715.594 | 86,87% | 3.715.585 | 86,87% |
| PVS2 | 12.367.809 | 9.338.953 | 75,51% | 8.676.335 | 92,90% | 8.676.312 | 92,90% |
| Egg1 | 21.504.101 | 10.673.452 | 49,63% | 9.219.560 | 86,38% | 9.219.503 | 86,38% |
| Egg2 | 12.581.422 | 10.673.452 | 84,84% | 10.202.002 | 95,58% | 10.201.718 | 95,58% |
| Emb1 | 40.816.766 | 36.273.048 | 88,87% | 33.723.248 | 92,97% | 33.721.943 | 92,97% |
| Emb2 | 35.811.987 | 32.006.382 | 89,37% | 29.818.243 | 93,16% | 29.817.043 | 93,16% |
| Nym1 | 32.934.837 | 30.502.999 | 92,62% | 29.155.402 | 95,58% | 29.155.221 | 95,58% |
| Nym2 | 27.271.760 | 25.385.570 | 93,08% | 24.245.468 | 95,51% | 24.245.380 | 95,51% |

#### Supplementary Table 2

| Sample | Number of reads associated with repeat elements | Number of piRNA reads | Number of Paired piRNA Reads | Number of Paired piRNA Reads (%) | Number of Unpaired piRNA Reads | Number of Unpaired piRNA Reads (%) |
|---|---|---|---|---|---|---|
| pvs1 | 869.438 | 138.187 | 17.284 | 12,51 | 120.903 | 87,49 |
| pvs2 | 2.011.369 | 405.222 | 56.006 | 13,82 | 349.216 | 86,18 |
| egg1 | 3.818.141 | 531.599 | 77.256 | 14,53 | 454.343 | 85,47 |
| egg2 | 5.003.045 | 3.873.579 | 714.790 | 18,45 | 3.158.789 | 81,55 |
| emb1 | 16.909.817 | 13.642.074 | 4.033.460 | 29,57 | 9.608.614 | 70,43 |
| emb2 | 15.678.915 | 13.054.204 | 3.836.780 | 29,39 | 9.217.424 | 70,61 |
| nym1 | 4.944.981 | 3.183.897 | 931.758 | 29,26 | 2.252.139 | 70,74 |
| nym2 | 2.825.846 | 1.230.413 | 341.111 | 27,72 | 889.302 | 72,28 |
| total | 52.061.552 | 36.059.175 | 10.008.445 | - | 26.050.730 | - |
